# The discovery of three new hare lagoviruses reveals unexplored viral diversity in this genus

**DOI:** 10.1101/466557

**Authors:** Jackie E. Mahar, Robyn N. Hall, Mang Shi, Roslyn Mourant, Nina Huang, Tanja Strive, Edward C. Holmes

## Abstract

Our knowledge of mammalian viruses has been strongly skewed toward those that cause disease in humans and animals. However, recent metagenomic studies indicate that most apparently healthy organisms carry viruses, and that these seemingly benign viruses may comprise the bulk of virus diversity. The bias toward studying viruses associated with overt disease is apparent in the lagoviruses (family *Caliciviridae*) that infect rabbits and hares: although most attention has been directed toward the highly pathogenic members of this genus - the rabbit haemorrhagic disease virus and European brown hare syndrome virus - a number of benign lagoviruses have also been identified. To determine whether wild European brown hares in Australia might also carry undetected benign viruses, we used a meta-transcriptomics approach to explore the gut and liver RNA viromes of these invasive animals. This led to the discovery of three new lagoviruses. While one of the three viruses was only detected in a single hare, the other two viruses were detected in 20% of all hares tested. All three viruses were most closely related to other hare lagoviruses, but were highly distinct from both known viruses and each other. We also found evidence for complex recombination events in these viruses, which, combined with their phylogenetic distribution, suggests that there is likely extensive unsampled diversity in this genus. Additional metagenomic studies of hares and other species are clearly needed to fill gaps in the lagovirus phylogeny and hence better understand the evolutionary history of this important group of mammalian viruses.

## Introduction

Although viruses probably infect all cellular organisms (1, 2), their true diversity is both underappreciated and poorly understood (3). Until recently, virus discovery was generally challenging, with a reliance on cell culture and PCR-based techniques. This, combined with a focus on viruses of anthropogenic importance, has meant that the great majority of viruses studied are those causing disease in humans and animals of human interest (2), generating a skewed perception of virus diversity and perhaps of virus evolution. Recent studies using next generation sequencing and metagenomics have demonstrated that a wealth of viruses exist in apparently healthy vertebrate and invertebrate organisms, and that the characterization of these viruses is vital to understanding virus evolution and ecology (4-6). The study of such “background” viruses in vertebrates is of particular interest in this context, as these can have the capacity to jump species boundaries and sometimes evolve new levels of virulence (7-10).

Lagoviruses are an example of a group of viruses within which both benign and highly pathogenic viruses have been detected (11-16). *Lagovirus* is a genus in the family *Caliciviridae*, comprising positive-sense, single stranded RNA viruses that infect members of the *Leporidae* family of mammals (i.e. rabbits and hares) (14, 17, 18). *Lagovirus* genomes are approximately 7.5 kb in length and are made up of two open-reading frames (ORF), one which encodes a polyprotein that is proteolytically cleaved to produce the non-structural proteins and the major capsid protein, VP60; and a second ORF that encodes the minor structural protein (17, 19, 20). Similar to other caliciviruses, the prototype lagoviruses have been shown to possess a subgenomic RNA, which is collinear with the 3’ end of the genomic RNA and encodes the structural genes (17, 21-24). Viruses in this genus have been classified into two proposed genogroups: GI encompasses all viruses related to rabbit haemorrhagic disease virus (RHDV, GI.1), which are generally rabbit-specific viruses, while GII includes viruses related to European brown hare syndrome virus (EBHSV, GII.1), most of which are hare-specific viruses (13). Pathogenic viruses of the *Lagovirus* genus, such as RHDV, RHDV2 (GI.2) and EBHSV, primarily affect the liver, causing massive hepatic necrosis usually associated with high mortality (25). In contrast, benign viruses exhibit an intestinal tropism (11, 12, 14).

Lagoviruses are a relatively well studied group of viruses since the high mortality rates of RHDV and RHDV*2* in the European rabbit (*Oryctolagus cuniculus*) have major ecological and economic impacts in Europe and Australia, although for different reasons (18). In parts of Europe, rabbits are an important part of the natural ecosystem, and are also farmed for rabbit meat and fur, upon which RHDV epidemics can have devastating effects (18). Conversely, rabbits are a pest species in Australia, and RHDV is used to control overabundant rabbit populations (18).

In Australia, extensive work has been done to characterise rabbit lagoviruses due to their importance in rabbit biocontrol. European rabbits and European brown hares (*Lepus europaeus*) were both successfully introduced into Australia by European settlers in the 1800s (26, 27), and both eventually reached plague densities, causing agricultural and ecological damage (28). While rabbits successfully colonized most of the continent except the wet tropics and extremely arid zones (29), hares did not spread as far, occupying a region of approximately 7 × 10^4^ km^2^ in size (26), primarily in the south-east of Australia. For a number of unconfirmed reasons, the hare population appeared to decline rapidly at the start of the 1900s (28), although they are still considered as pests in some areas of Australia today (30, 31). Rabbits became a major pest species, and accordingly, RHDV was deliberately introduced into Australia as a biocontrol agent in the mid-1990s, and continues to be released periodically (32).

RHDV is not the only lagovirus present in Australia. After the initial release, reduced effectiveness of RHDV was noted in south-eastern temperate regions of Australia, and serological data suggested the existence of a related benign virus (33). Subsequently, a benign lagovirus, RCV-A_1_ (GI.4) was isolated and sequenced (14), and shown to confer a degree of cross-protection against RHDV, potentially interfering with biocontrol (34, 35). Additionally, a number of seemingly benign lagoviruses were detected in Europe and New Zealand, some closely related to RCV-A1 and others more closely related to pathogenic lagoviruses (11-13, 36, 37). In the last decade, a number of other lagoviruses have appeared in Australia, including RHDVa-Aus (GI.4eP-GI.1a) (38), RHDV2 (GI.2) (39) and a recombinant of these two variants (GI.4eP-GI.2, [RdRp-capsid genotype]) (13, 40). RHDV_2_ has also been detected in hares (*Lepus europaeus*) in Australia (41), and although phylogenetic analysis suggests that these infections were likely the result of spill-over from sympatric rabbit populations, this virus is highly virulent in hares (42). Apart from RHDV_2_, no other lagoviruses have been detected in hares in Australia. In Europe, *Lepus europaeus* is affected by the pathogenic lagovirus EBHSV (43-48), which has never been detected in Australia, as well as RHDV2, which has been detected in multiple hare species (41, 49-52). Two additional, presumably benign, hare caliciviruses from Europe (denoted GII.2), have recently been reported (13, 15, 16) and it is unknown whether similar viruses are present in Australia.

Less emphasis has been placed on the investigation of viruses in hares compared to those affecting rabbits, possibly because unlike rabbits, hares are not commercially farmed for meat and have a comparatively lower impact as a pest species in Australia. However, exploration of the hare virome is of importance not just for understanding the biological and genetic diversity of lagoviruses, but also for understanding the frequency with which these viruses have been able to change their virulence and host range within lagomorphs throughout the evolutionary history of this genus. We therefore aimed to explore the RNA virome of healthy hares in Australia to detect and characterize unidentified viruses, and in doing so, broaden our understanding of RNA virus diversity and evolution.

## Results

### Initial PCR for lagovirus detection

As an initial screening method to identify samples likely to contain diverse lagoviruses, a broad-range universal lagovirus PCR (14) was used to analyse hare duodenum samples (n = 38) collected in two locations, Hamilton, Victoria (VIC) and Mulligan’s Flat, Australian Capital Territory (ACT). Lagoviruses were detected in two hare duodenum samples: MF-150 and JM-2. This was confirmed by Sanger sequencing, and initial phylogenetic analysis of the ~300 nt sequences indicated that these viruses were distinct from known lagoviruses and each other (data not shown).

### RNA Sequencing

In an effort to obtain the complete genome of the new lagoviruses, RNA sequencing (i.e. “meta-transcriptomics” (5)) was performed on JM-2 and MF-150 duodenum RNA and a selection of other hare duodenum and liver RNA pooled into 16 libraries (Table 1). An aggregate of 792,023,978 reads were obtained for all libraries, 605,624,790 (76%) of which did not map to host rRNA, averaging 38,086,664 non-rRNA reads per library.

**Table 1.** Prevalence of new viruses and sequencing library details.

| Name | Sample collection |  | Sequencing library<sup>†</sup> |  | Virus detection by PCR in duo<sup>‡</sup> |  |  |
| --- | --- | --- | --- | --- | --- | --- | --- |
|  | Location* | Date | Liver | Duodenum | HaCV-A1 | HaCV-A2 | HaCV-A3 |
| JM-1 | Ham | 30/6/16 | N/A | N/A | - | - | - |
| JM-2 | Ham | 30/6/16 | N/A | JM-2-duo | + | - | - |
| JM-3 | Ham | 30/6/16 | N/A | N/A | - | - | - |
| JM-4 | Ham | 30/6/16 | N/A | N/A | - | + | - |
| JM-5 | Ham | 30/6/16 | N/A | N/A | - | - | - |
| JM-6 | Ham | 30/6/16 | N/A | N/A | - | + | - |
| JM-7 | Ham | 30/6/16 | N/A | N/A | - | - | - |
| JM-8 | Ham | 30/6/16 | N/A | N/A | - | - | - |
| JM-9 | Ham | 30/6/16 | N/A | N/A | - | + | - |
| JM-10 | Ham | 30/6/16 | N/A | N/A | - | - | - |
| JM-11 | Ham | 30/6/16 | N/A | N/A | - | +(L) | - |
| JM-12 | Ham | 30/6/16 | N/A | N/A | - | + | - |
| JM-13 | Ham | 30/6/16 | N/A | N/A | - | - | - |
| JM-14 | Ham | 30/6/16 | N/A | N/A | - | - | - |
| JM-15 | Ham | 30/6/16 | N/A | N/A | - | - | - |
| JM-16 | Ham | 30/6/16 | N/A | N/A | + | - | - |
| JM-17 | Ham | 30/6/16 | N/A | N/A | - | - | - |
| JM-18 | Ham | 30/6/16 | N/A | N/A | - | - | - |
| JM-19 | Ham | 30/6/16 | N/A | N/A | - | - | - |
| JM-20 | Ham | 30/6/16 | N/A | N/A | - | + | - |
| JM-22 | Ham | 23/5/17 | Ham1-L | Ham1-D | + | - | - |
| JM-24 | Ham | 23/5/17 | Ham1-L | Ham1-D | + | + | - |
| JM-26 | Ham | 23/5/17 | Ham1-L | Ham1-D | - | +/- | - |
| JM-27 | Ham | 23/5/17 | Ham2-L | Ham2-D | +/- | - | - |
| JM-29 | Ham | 23/5/17 | Ham2-L | Ham2-D | + | - | - |
| JM-30 | Ham | 23/5/17 | Ham3-L | Ham3-D | + | - | - |
| JM-31 | Ham | 23/5/17 | Ham3-L | Ham3-D | - | - | - |
| JM-34 | Ham | 23/5/17 | Ham3-L | Ham3-D | - | +(L) | - |
| JM-35 | Ham | 23/5/17 | Ham4-L | Ham4-D | + | - | - |
| JM-40 | Ham | 23/5/17 | Ham4-L | Ham4-D | +(L) | - | - |
| MF-01 | MF | 20/12/12 | N/A | N/A | - | - | - |
| MF-02 | MF | 20/12/12 | N/A | N/A | - | + | - |
| MF-07 | MF | 20/12/12 | MF1-L | MF1-D | - | - | - |
| MF-22 | MF | 20/12/12 | MF1-L | MF1-D | - | - | - |
| MF-137 | MF | 3/2/16 | MF1-L | MF1-D | - | - | - |
| MF-148 | MF | 5/5/16 | MF2-L | MF2-D | - | - | - |
| MF-149 | MF | 5/5/16 | MF2-L | MF2-D | - | - | - |
| MF-150 | MF | 9/6/16 | MF3-L | MF3-D, MF-150 | - | - | + |
| MF-151 | MF | 9/6/16 | MF3-L | MF3-D | - | - | - |
| MF-152 | MF | 9/6/16 | MF3-L | MF3-D | - | - | - |
| MF-155 | MF | 7/7/16 | MF4-L | MF4-D | - | - | - |
| MF-156 | MF | 27/7/16 | MF4-L | MF4-D | - | - | - |

\*Ham, Hamilton Victoria; MF, Mulligan's Flat, Australian Capital Territory.
<sup>†</sup>N/A, not applicable – RNA sequencing was not performed on these samples703 <sup>‡</sup>duo, duodenum; +, positive; -, negative; +/-, weak positive; (L) weak positive in liver RNA

Reads were assembled into contigs and screened for viruses. No viral contigs were assembled from any of the liver libraries or most of the Mulligan’s Flat duodenum libraries. The four Hamilton duodenum pooled libraries, one Mulligan’s Flat duodenum pooled library (MF3-D), and the JM-2 duodenum library together had a total of 58 viral contigs (Figure 1), which matched seven different viruses in a BLAST analysis. Contigs with highest identity to lagoviruses (either EBHSV [GII.1] or hare calicivirus [GII.2]) were present in all of these libraries and, on average, made up 86% (71–100%) of all viral contigs in each library (Figure 1). However, lagovirus reads were in very low abundance overall, comprising less than 0.003% of the non-rRNA transcriptome in each library. The 50 lagovirus contigs had an average nucleotide identity of only 84.7–89.9% to the top BLAST result, indicating a potential new virus. The longest of the lagovirus contigs (5,586 nt) encompassed ~75% of a typical lagovirus genome, while the shortest was 201 nt.

**Figure 1.**
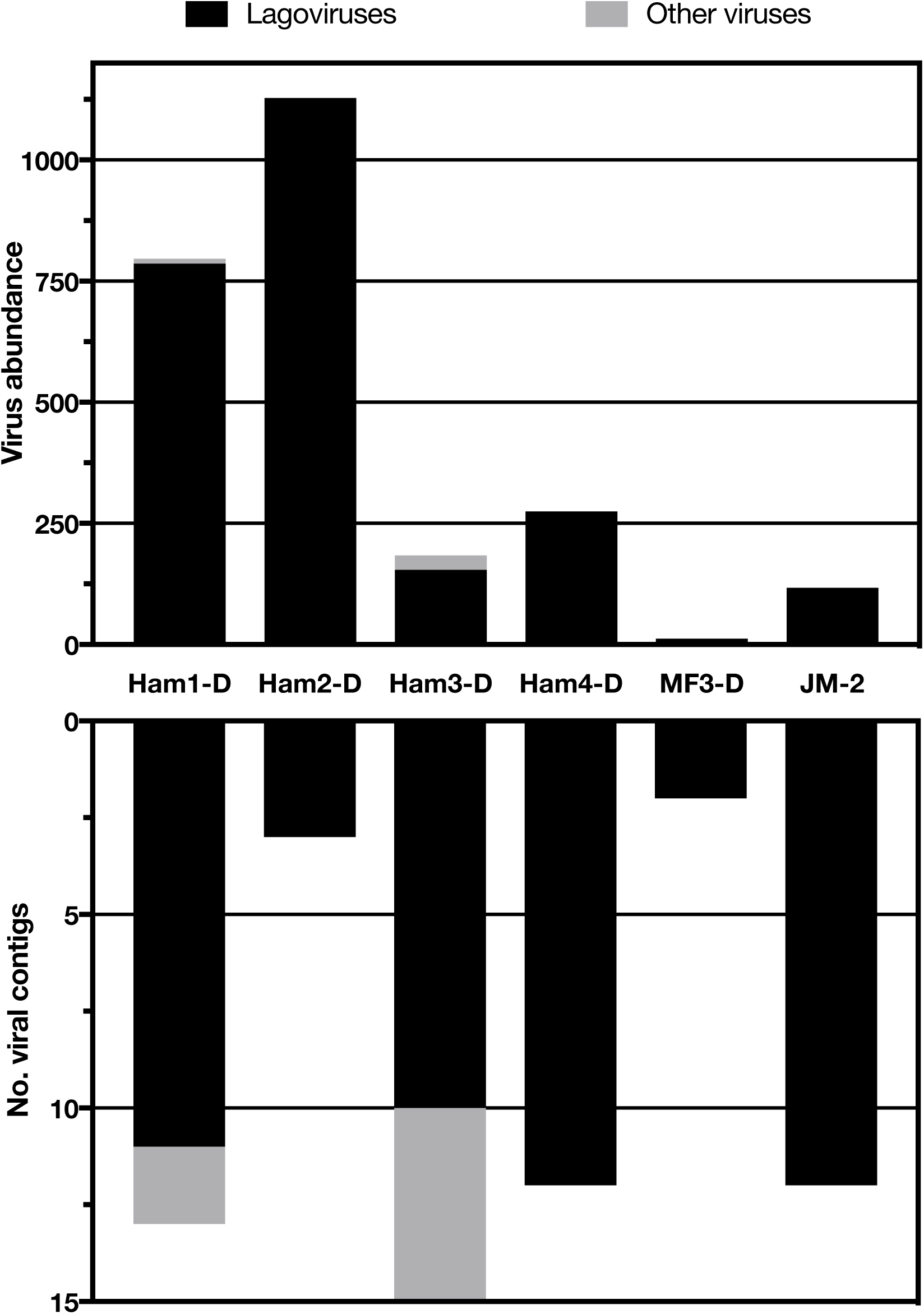
Number of virus contigs and viral abundance in hare duodenum libraries. Top - the relative abundance (reads, expected count) of viruses (y-axis) in each hare duodenum library (x-axis). Bottom - the number of viral contigs (y-axis) that were assembled for each library (x-axis). Libraries for which no virus contigs were generated are not shown. The bars are shaded according to virus type where black represents lagovirus contigs and grey represents all other virus genera. Ham, Hamilton library; MF, Mulligan’s Flat library; JM-2, specific sample from Hamilton.

Apart from lagoviruses, contigs were assembled for four other putative viruses, with the closest BLAST hits to: Hubei partiti-like virus 54, Hubei partiti-like virus 49, Mammalian orthoreovirus 1 and Mammalian orthoreovirus 3. However, only short contigs (range 214–483 nt) were assembled for these viruses, and all had very low abundance of less than 20 reads (Figure 1).

### Lagovirus genome assembly and annotation

#### Hamilton hare viruses

Lagovirus contigs from the Hamilton libraries were assembled *de novo* to form “reference assemblies” and reads from individual libraries were mapped back to the reference assembly consensus sequences. Through this approach, an almost complete genome sequence was assembled for two new lagoviruses. The first, provisionally named Hare calicivirus Australia-1 (HaCV-A1) was assembled from the Ham-2D library and was 7,364 nt in length with 22.8X mean coverage. The second virus comprised two assemblies (5,588 nt and 1,628 nt) that did not overlap, but were later shown to be from the same virus using PCR and Sanger sequencing. This virus was provisionally named Hare calicivirus Australia-2 (HaCV-A2) and both assemblies for this virus were generated from the Ham-1D library with a mean coverage of 19.3X and 10.9X.

To validate our RNA sequencing and genome assembly approach, we confirmed the HaCV-A1 genome sequence by amplicon sequencing of an individual sample. There were only ten nucleotide differences between the amplicon-based sequencing approach and the *de novo* RNA sequencing assembly approach, and these occurred at highly variable sites (data not shown). This level of variation is to be expected given that amplicon sequencing was conducted on a single sample while, in contrast, the RNA sequencing was conducted on a pool of samples. As part of the amplicon-based sequencing approach, we were also able to determine the 3’ end of the virus, revealing a 64 nt 3’ UTR. The start of the coding sequence, as well as 3 nt of the 5’ UTR, was determined at the 5’ end, although the first 8 nt of the 5’ end sequence are inferred as these were obtained from amplicon sequencing (only) and are within the primer binding region. Excluding the 5’ end primer inferred sequence, and the polyA tail, a consensus sequence of 7,386 nt was obtained for HaCV-A1 (Figure 2).

**Figure 2.**
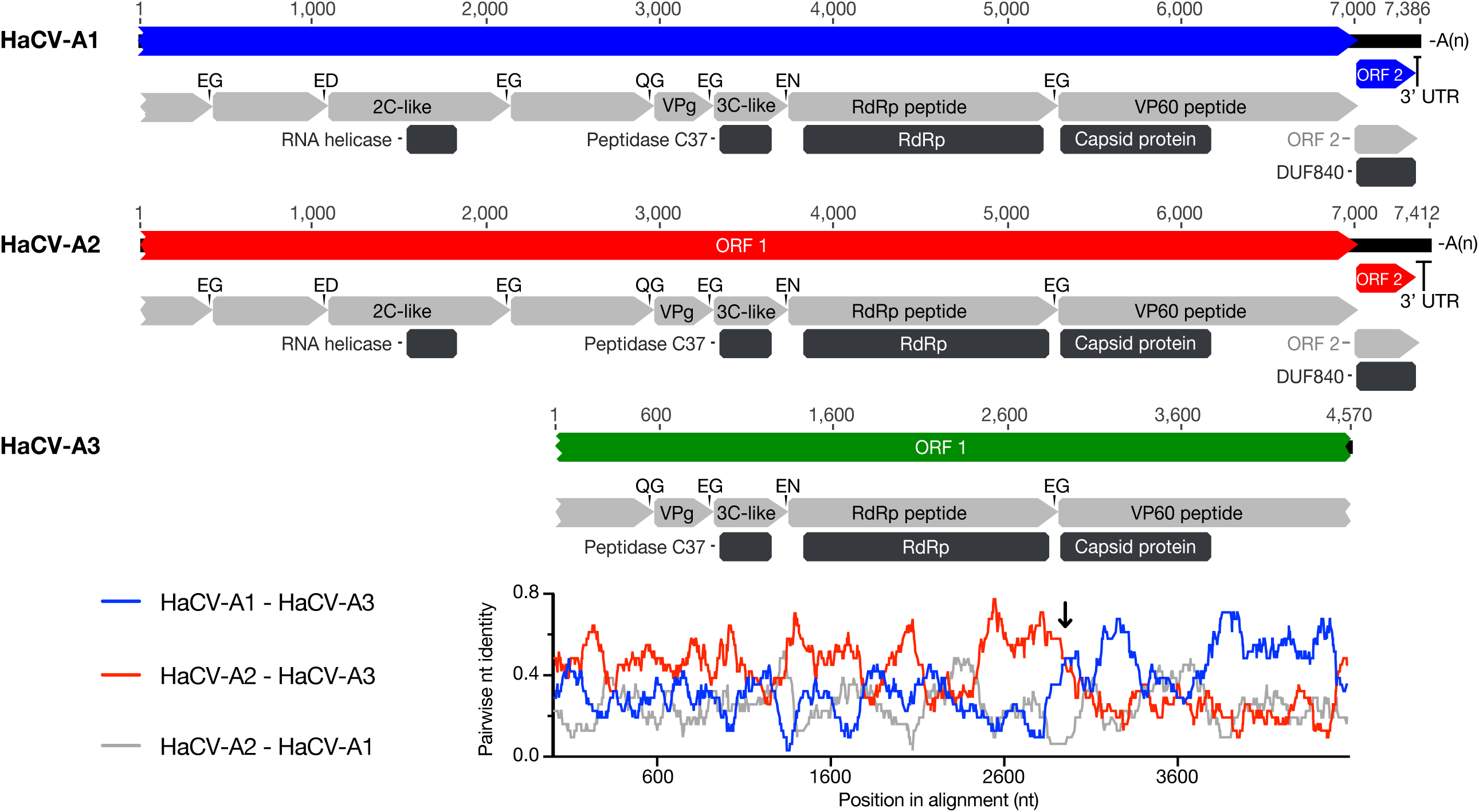
Genome structure of new viruses and identity plot. A schematic representation of the region of the genome sequenced for each new virus is shown above a pairwise identity plot. Open-reading frames (ORFs) are represented by coloured arrow bars (blue, HaCV-A1; red, HaCV-A2; green, HaCV-A3). Conserved protein domains detected using the NCBI conserved domains search tool, are indicated by dark grey boxes (RdRp, RNA-dependent RNA polymerase; DUF840, lagovirus protein of unknown function). The likely cleavage fragments/peptides of the ORF1 polyprotein, inferred from sequence homology with EBHSV and RHDV, are indicated by the light grey arrow bars (2C-like, 2C-like RNA helicase; VPg, genome-linked viral protein; 3C-like, 3C-like proteinase; RdRp, RNA-dependent RNA polymerase; VP60, major capsid protein). Amino acids flanking the likely cleavage sites in the polyprotein are indicated at the junction between the peptides using amino acid l-letter identifiers. Note that a broken appearance at either end of the arrow bars indicates incomplete sequence for that ORF or peptide. The 3’ untranslated regions (UTR) and polyA tails (A(n)) are indicated where sequence was obtained. Genome numbering at regular intervals is indicated above each schematic. Pairwise nucleotide identity (y-axis) according to genome position (x-axis) is plotted below the HaCV-A3 schematic. The plot was generated in the RDP4 program from an alignment trimmed to the length of HaCV-A3, using a sliding window of 30 nt. A clear cross-over between the blue line (identity between HaCV-A3 and HaCV-A1) and the red line (identity between HaCV-A3 and HaCV-A2) suggests that HaCV-A3 is a recombinant between parental viruses related to HaCV-A1 and HaCV-A2. The cross-over event occurs at the junction of the RdRp and capsid and is indicated by a black arrow.

Additional amplification and sequencing was also conducted for HaCV-A2 to bridge the gap between the two assemblies and join them together, to confirm regions with gaps or low coverage, and to extend the sequence. This resulted in a consensus sequence of 7,412 nt, including a 3’ UTR of 77 nt (Figure 2). The complete 5’ end was not obtained, with 14 nt likely missing from the start of ORF1 (based on sequence similarity with other lagoviruses).

Both HaCV-A1 and HaCV-A2 appear to have the same genome organisation as other lagoviruses, with two ORFs: one encoding a polyprotein containing the non-structural genes and capsid gene, and one encoding the minor structural protein (Figure 2). The polyprotein encoded by ORF1 is likely 2,332 amino acids in length for HaCV-A1 (based on start codon in the primer inferred sequence) and is likely the same for HaCV-A2, although the start of the coding sequence was not obtained for the latter virus. The likely cleavage products (mature peptides) resulting from post-translational processing of the ORF 1 polyprotein (and cleavage sites) were inferred from sequence similarity with EBHSV and RFIDV. These included peptides for which conserved domains were identified (RNA helicase, peptidase C37/3C-like proteinase, RNA-dependent RNA polymerase (RdRp), calicivirus capsid protein, and DUF840, a lagovirus protein of unknown function), as well as the genome-linked viral protein, VPg, which binds to the 5’ end of calicivirus RNA molecules (21); and three proteins with unknown function, as indicated in Figure 2. Potential termination upstream ribosomal binding site (TURBS) motifs were identified at positions 6,904-6,908 (motif 2^⋆^), 6,910-6,916 (motif 1), 6,959-6,963 (motif 2) of the HaCV-A1 partial genome sequence and at the equivalent location in the HaCV-A2 partial genome sequence; 6,907-6,911 (motif 2^⋆^), 6,913-6,919 (motif 1), and 6,962-6,966 (motif 2). For both viruses, based on the putative location of the TURBS motifs (53), the second ORF is likely to overlap with the first ORF by 8 nt, as seen for EBHSV (17), and encode a protein of 113 amino acids.

#### Mulligan’s Flat hare virus

Only two short (~200 nt) lagovirus contigs were assembled from the Mulligan’s flat libraries using a meta-transcriptomics approach. Flowever, almost the complete capsid gene (1,589 nt) of the lagovirus detected in MF-150 duodenum was amplified and Sanger sequenced using a combination of primers for the EBHSV capsid protein gene (54) and specifically designed broadly-reactive primer sets. These Sanger sequences, together with reads from RNA sequencing and the two contigs from the Mulligan’s Flat libraries, were mapped to an EBHSV reference sequence. Six reads and the two contigs mapped to regions of the genome upstream from the Sanger-sequenced capsid protein gene, enabling further amplification of the intervening regions. Subsequent amplicon sequencing extended the sequenced region of this virus to 4,570 nt (Figure 2). This virus is distinct from HaCV-A1 and HaCV-A2 and was provisionally named Flare calicivirus Australia-3 (HaCV-A3). Although the complete genome was not isolated for HaCV-A3, the genome organisation of the obtained sequence appears to match that of known lagoviruses (Figure 2). Conserved domains for peptidase C37/3C-like proteinase, RdRp, and calicivirus capsid protein were identified, as well as likely cleavage products of the ORF1 polyprotein, including VPg, 3C-like proteinase, RdRp, capsid protein and part of an unknown protein at the 5’ end of the sequence (Figure 2).

### Phylogenetic analysis

The three new hare lagoviruses were diverse, with HaCV-A1 sharing only 74% nt and 77% nt identity across sequenced regions with HaCV-A2 and HaCV-A3, respectively; while HaCV-A2 and HaCV-A3 share 78% nt identity. Notably, HaCV-A3 was most similar to HaCV-A2 in the RdRp gene and most similar to HaCV-A1 in the capsid region (Figure 3), suggestive of recombination.

**Figure 3.**
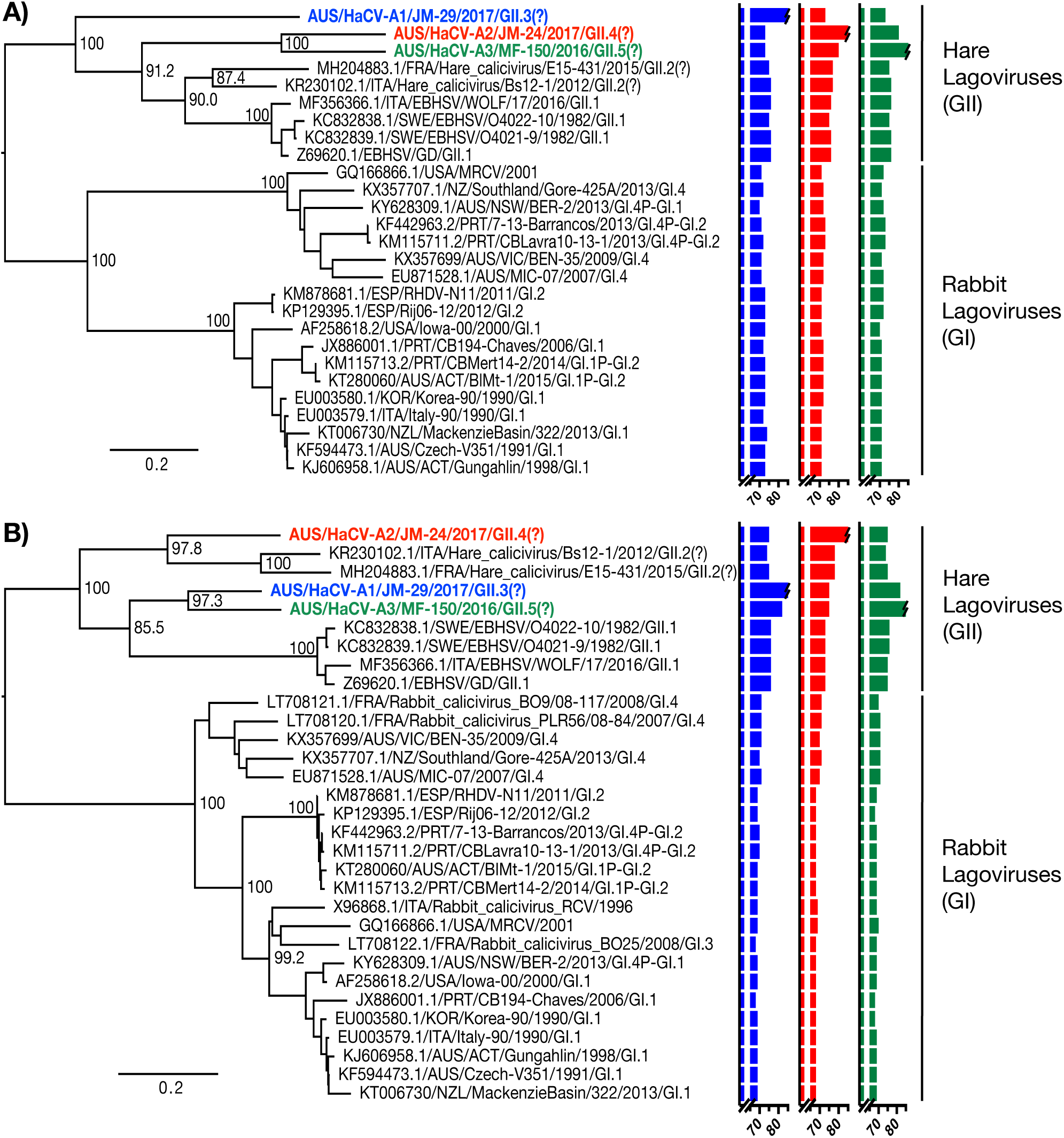
Phylogenetic analysis of new lagovi ruses. Maximum likelihood phylogenies of the (A) RdRp gene (n=27; 1,548 nt) and (B) capsid gene (n=3i; 1,704 nt) were inferred for the three new lagoviruses along with representative members of the genus *Lagovirus.* The accession number of sequences obtained from GenBank is shown in the taxa labels. Trees were mid-point rooted for clarity only, and branch support was estimated using 1,000 bootstrap replicates, which are shown at the major nodes. The taxa names of the three benign hare viruses reported in this study are coloured blue (HaCV-A1), red (HaCV-A2), and green (HaCV-A3). Graphs to the right of the trees indicate the percentage nucleotide identity (x-axis) of each of the three new viruses (HaCV-A1, blue; HaCV-A2, red; HaCV-A3, green) with other taxa in the trees for both genes. The two major clades in each phylogeny are labelled according to proposed genogroup and prototypical host. Proposed genotypes are indicated in taxa labels (those without a genotype label are unclassified).

Maximum likelihood phylogenetic trees were inferred for both the RdRp and capsid genes of the new viruses together with representative lagoviruses (Figure 3). The three new hare lagoviruses form a strongly supported monophyletic group with other hare-specific lagoviruses (G1l) – EBHSV and hare caliciviruses – in both the capsid and RdRp gene phylogenies. Flowever, all three viruses are clearly distinct, separated both from each other and other known lagoviruses by relatively long branches with strong bootstrap support (Figure 3). Indeed, there is approximately the same phylogenetic distance between HaCV-A1 and HaCV-A2 as among all known rabbit lagoviruses. Given this diversity, and according to the proposed classification guidelines outlined by Le Pendu et al (2017), it is likely that each of these viruses would represent a new genotype within the GII genogroup of the *Lagovirus* genus, provisionally GII.3 (HaCV-A1), GII.4 (HaCV-A2) and GII.5 (HaCV-A3). A substantial level of phylogenetic incongruence between the RdRp and capsid protein gene trees is evident among the hare caliciviruses (Australian and European). In the capsid gene tree (Figure 3B), HaCV-A2 clusters most closely with the two European benign hare caliciviruses (GII.2), sharing 79% and 78% nt identity with each, while the other two new viruses cluster together and share a more recent common ancestor with the pathogenic hare lagovirus, EBHSV. In the RdRp phylogeny (Figure 3A), the two European hare caliciviruses cluster most closely with EBHSV and the three viruses discovered here are more distant.

### Lagovirus recombination

As our phylogenetic analysis strongly suggested the occurrence of recombination among the new viruses, particularly the incongruence between the RdRp and capsid gene phylogenies, we performed a more detailed analysis of this putative recombination event. These analyses revealed strong evidence of recombination among the Australian hare caliciviruses. Specifically, HaCV-A3 was predicted to be a recombinant of viruses related to HaCV-A1 and HaCV-A2 (RDP pairwise distance plot; Figure 2). The estimated location of the putative breakpoint was at position 2,909 in the HaCV-A3 sequence (99% CI: 2,827–3,183), 19 nucleotides downstream of the RdRp/capsid putative cleavage site, which is the equivalent of position 5,307 on the reference EBHSV genome sequence (accession NC_002615). Importantly, recombination at this location essentially divides the genome into a region encoding the non-structural proteins and a second region encoding the structural proteins (Figure 2). Phylogenetic analysis on regions either side of the breakpoint strongly supported the occurrence of recombination, with HaCV-A3 clustering with HaCV-A2 in the non-structural genes tree (left of the breakpoint) and clustering with HaCV-A1 in the capsid tree (right of the breakpoint), with robust bootstrap support (>70%). This phylogenetic incongruence is captured in the RdRp and capsid phylogenies presented in Figure 3. However, an end breakpoint could not be determined. Given the substantial diversity of potential parent sequences and lack of sampling inthisclade, it was difficult to predict the evolutionary history of recombinant events with certainty. Accordingly, HaCV-A1 or HaCV-A2 may be the actual recombinant, or it is possible that more than one recombination event has occurred among these and related viruses. Notably, the regions flanking the putative recombination breakpoint were amplified in a single amplicon for each of the Australian hare caliciviruses, excluding the possibility of miss-assembly leading to false recombination signals.

### Prevalence of new hare lagoviruses

Specific screening PCRs amplifying a short region of the capsid gene were designed for each new virus, and 42 duodenum samples from shot healthy hares from both locations were tested for the presence of these viruses (including those used to generate the sequencing libraries). Accordingly, HaCV-A1 and HaCV-A2 were both found at a prevalence of 30% in Hamilton, VIC, and one hare was infected with both (Table 1). HaCV-A1 was not detected in Mulligan’s Flat, ACT, while HaCV-A2 was detected in 1/12 rabbits in this location, making it the only virus of the three newly identified lagoviruses to be detected at both locations (Table 1). HaCV-A3 was detected in only one hare duodenum, MF-150, and was likely to be present at a very low concentration, as viral contigs could not be assembled from a total of 39,993,840 reads from RNA sequencing of this sample. For PCR-positive duodenum samples, liver samples from the same individuals were also screened for the presence of the new lagoviruses. HaCV-A1 was detected in one liver sample and HaCV-A2 was detected in two liver samples (Table 1), although the amplicons were faint, indicating a lower viral abundance compared to the duodenum or possible contamination during sample collection.

## Discussion

We used a bulk RNA-Sequencing approach to explore the RNA virome of European brown hares in Australia. This resulted in the discovery of three new hare viruses: Hare calicivirus Australia-1 (HaCV-A1), Hare calicivirus Australia-2 (HaCV-A2) and Hare calicivirus Australia-3 (HaCV-A3). Prior to this, the only lagovirus detected in hares in Australia was RHDV2 (41), which is primarily a rabbit virus, and phylogenetic evidence suggests that RHDV2 infection in hares in Australia occurred as a result of transient spill-over events (42, 49). While HaCV-A3 was only found in one animal, the other two new viruses were both detected in almost one third of hares tested at the Hamilton, VIC site, during both sampling periods (one year apart). This suggests that similar to the non-pathogenic rabbit calicivirus RCV-A1 (GI.4) (55), these two viruses may be prevalent in certain populations, although more extensive screening is needed to confirm this.

The three new viruses were all members of the genus *Lagovirus* (family *Caliciviridae*). This genus comprises both virulent and benign viruses that infect hares (*Lepus*) and rabbits (*Oryctolagus cuniculus*) (14). The virulent viruses, RHDV (GI.1), RHDV2 (GI.2), and EBHSV (GII.1) have a liver tropism and are associated with necrotic hepatitis often resulting in fatality (25, 56, 57), while the benign viruses, RCV-A1 (GI.4), RCV (unclassified), RCV-E1 (G1.3), and the French hare calicivirus (G1l.2), have an intestinal tropism (11, 12, 14, 16). Tissue tropism has not been reported for the Italian benign hare calicivirus (15). All three new viruses discovered here were found in low abundance in the duodenum of apparently healthy hares, consistent with the intestinal tropism observed for benign lagoviruses (11, 14, 16). Either HaCV-A1 or HaCV-A2 were also detected in the liver of three of these apparently healthy hares, although at objectively lower levels, suggesting that the site of replication is likely to be the intestine. The genome organization of the new viruses was consistent with that of other lagoviruses, and the putative proteolytic cleavage sites on the ORF1 polyprotein are identical to those in EBHSV, suggesting similar processing mechanisms in these new viruses (19).

Notably, there was evidence of recombination between the three new viruses, confirming the importance of recombination as a means to generate genetic diversity in lagoviruses (40, 58). The location of the putative breakpoint is near the junction of the RdRp and capsid protein genes. Recombination in this region results in chimeric viruses with non-structural genes derived from one virus and structural genes derived from another (22, 23, 40, 59). This is a recombination hotspot in caliciviruses, and similar events have been observed between members of the GI lineage of lagoviruses, where several recombinants between RHDV (GI.1), RHDV2 (GI.2) and RCV-A1(G1.4) have been reported (40, 58). The sequence in the RdRp/capsid junction is highly conserved in caliciviruses, and is predicted to form a stem loop structure that may facilitate a pause in replication and a subsequent template switch (22, 23). In addition, several caliciviruses have been shown to possess a subgenomic RNA encoding the structural genes, which may serve as an ideal secondary template for reinitiation of RNA synthesis (21-24). Due to the diversity between the three new Australian hare caliciviruses and related European hare caliciviruses, and general under-sampling of the *Lagovirus* GII clade, it is difficult to establish which of the new viruses is the recombinant and which is the parent, although our analysis suggested that HaCV-A3 was the most likely recombinant. Indeed, the pattern of phylogenetic incongruence between the RdRp and capsid gene trees may mean that several recombination events have occurred among these and related strains, although this may be difficult to detect/confirm due to under-sampling of potential parental sequences, because they have been over-written by more recent recombination events, or substantial divergence since putative recombination events (60).

The viruses identified here were most closely related to the only two known hare-specific lagoviruses (G1l), EBHSV (GII.1) and the recently reported European hare caliciviruses (denoted G1l.2) (13, 15, 16). The three new viruses are strikingly distant from the previously characterized hare lagoviruses and from each other, and each would constitute a new genotype of the *Lagovirus* genus (13), provisionally G1l.3 (HaCV-A1), G1l.4 (HaCV-A2) and G1l.5 (HaCV-A3). The addition of these three viruses to the lagovirus genus has therefore greatly increased the phylogenetic depth and diversity of this genus, indicating that lagoviruses have likely circulated for longer than previously assumed. The relatively large genetic distance both among these viruses and between the hare and rabbit viruses almost certainly reflects a lack of sampling, with the possibility that lagoviruses may in fact infect a more diverse range of mammalian taxa, such as *Sylvilagus sp.* It should be noted that the two European hare caliciviruses, both tentatively denoted GII.2 (13,15), cluster together, but probably exhibit enough diversity to be classified as two different genotypes, with 85% nucleotide identity in the capsid protein gene (Figure 3).

The lagovirus genus is of evolutionary interest as virulence has likely evolved independently in RHDV (GI.1), RHDV2 (G1.2) and EBHSV (GII.1) (61). Historically, research efforts have been focused towards these highly virulent viruses due to their apparent impacts, however, non-virulent lagoviruses are increasingly being reported (11-14, 37). With additional comprehensive sequencing studies, the *Lagovirus* genus may indeed prove to be comprised of mainly asymptomatic viruses. These viruses may provide an extensive gene pool for recombination or even the possible emergence of additional virulent lagoviruses like RHDV2. Our data suggests that the *Lagovirus* genus is likely substantially undersampled and hence that the true diversity of the hare (and rabbit) virome is underestimated. Accordingly, extensive additional sequencing of RNA viromes of hares and other lagomorph species is needed to fill in the gaps in the lagovirus phylogeny and those of other RNA viruses, in turn providing broad-scale insights into RNA virus evolution and ecology.

## Materials and methods

### Sample collection

Liver and duodenum were collected post-mortem from apparently healthy hares shot from a vehicle using a 0.22 calibre rifle, and frozen at −20°C. Samples were taken from 30 hares in Hamilton, VIC over two nights–30/06/2016 and 23/05/2017–and 12 hares from Mulligan’s Flat, ACT in December 2012, February 2016, or May - July 2016 (Table 1). Samples were collected as part of a routine vertebrate pest control program and lagovirus serological surveillance studies. All work was carried out according to the Australian Code for the Care and Use of Animals for Scientific Purposes with approval from the institutional animal ethics committee (ESAEC12-15 and CLWA16-02).

### RNA isolation

RNA was extracted from 20-30 mg of tissue using the Maxwell 16 LEV simplyRNA tissue kit and extraction robot (Promega, WI, USA) as per the manufacturer’s instructions.

### cDNA synthesis and PCR for detection of diverse lagoviruses

First-strand cDNA was prepared using Invitrogen Superscript^™^ IV Reverse Transcriptase (Thermofisher Scientific, MA, USA) according to the manufacturer’s instructions using 5 μ1 of RNA and 500 ng of Oligo(dT)(18mer) or 10 μM CaVuniR specific primer (Supplementary table S1). For cDNA prepared for 3’ end amplification, 10 μM of primer GV270 (62) was used.

Duodenum samples (n = 38) were screened for the presence of lagoviruses using a universal lagovirus PCR as described previously (14), and positive amplicons were confirmed by Sanger sequencing at the Australian Cancer Research Foundation (ACRF) Biomolecular Resource Facility (BRF) in Canberra, ACT.

### Initial amplification of HaCV-A3 for Sanger sequencing

Regions of the genome of HaCV-A3 (sample MF-150) were initially amplified using EBHSV primers EBHSV_VP6o_o1728R and EBHSV_VP6o_o813F (54), or specifically designed broadly reactive primers (Supplementary table S1). PCRs were conducted using Invitrogen Platinum *Taq* Polymerase High Fidelity kit (Thermofisher Scientific, MA, USA) according to the manufacturer’s protocol, using 1.5 μ1 of diluted cDNA (1:2) as template in a 25 μ1 reaction. Positive amplicons were sequenced at ACRF-BRF.

### RNA sequencing

#### RNA library construction and sequencing

RNA from two hare duodenum samples that tested positive in the lagovirus PCR (MF-150 and JM-2), as well as RNA from the liver and duodenum of 10 additional hares from Hamilton VIC, and 10 hares (including MF-150) from Mulligans Flat ACT, were selected for sequencing (Table 1). Freshly extracted RNA was treated using Invitrogen TURBO DNase (Thermofisher Scientific, MA, USA) and further purified and concentrated using the RNeasy MinElute cleanup kit (Qiagen, Hilden, Germany). RNA was quantified using the Qubit RNA Invitrogen Broad-range Assay kit with the Qubit Fluorometer v3.0 (Thermofisher Scientific, MA, USA), and further quantified and assessed for quality using the Agilent RNA 6000 nano kit and Agilent 2100 Bioanalyzer (Agilent Technologies, CA, USA). JM-2 and MF-150 duodenum RNA were each submitted for sequencing as a single library, while the remaining RNA samples were pooled in equal proportions by location and tissue type into pools of 2-3 individuals, totalling 18 libraries (Table 1). MF-150 duodenum RNA was sequenced individually, and was also included in the Mulligan’s Flat duodenum pools, as the virus loads initially detected in MF-150 duodenum RNA appeared to have been very low. Library preparation and sequencing was carried out at the Australian Genome Research Facility (AGRF, Melbourne) using the TruSeq total RNA library preparation kit (Illumina, CA, USA) with host rRNA depletion using the Illumina Ribo-Zero-Gold rRNA removal kit (Epidemiology). Paired-end sequencing (100 bp) was performed on the HiSeq 2500 sequencing platform.

#### Contig assembly and annotation

Reads were trimmed using Trimmomatic (63) and assembled into contigs *de novo* using Trinity (64). Abundance (as expected counts) was estimated for each contig using the RSEM tool (65), an alignment-based quantification method implemented in Trinity. BLASTn and DIAMOND BLAST× were used to compare Trinity contigs to the NCBI nucleotide (nt) database (e-value cut-off 1×10^10^) and non-redundant protein (nr) database (e-value cut-off I×10^5^), respectively. Results were filtered and contigs that had a viral hit for either BLAST search were retained. Virus host associations were allocated using the Virus-Host database (https://www.genome.jp/virushostdb). All reads were mapped to host rRNA (rabbit rRNA sequences were used as hare rRNA sequences were not available) using bowtie2 (66), to quantify remaining host rRNA reads since the laboratory-based steps are usually not sufficient to completely eliminate host rRNAs. The rabbit host rRNA target index was generated from a complete *O. cuniculus* 18s rRNA reference sequence obtained from GenBank (accession NR_033238) and a near complete *O. cuniculus* 28s rRNA sequence obtained from the Silva high quality ribosomal database (67) (accession GBCA01000314). The total number of reads that were not mapped to host rRNA for each library were used as the denominator to calculate the percentage of reads mapped to viral contigs.

#### Lagovirus genome assemblies

To increase the chance of assembling entire viral genomes, the lagovirus contigs from all libraries, in addition to Sanger sequences obtained for JM-2 and MF-150 duodenum samples, were aligned, and contigs with overlapping regions were merged, using the Geneious assembler (68) (with the highest sensitivity setting). Four merged contigs were generated with lengths 7,364 nt, 5,588 nt, 1,628 nt, and 1,375 nt. The three longest merged contigs were generated from contigs from the Hamilton, VIC libraries, while the shortest was compiled from Sanger sequences from MF-150 duodenum (Mulligan’s Flat, ACT). To generate library-specific lagovirus contigs, the consensus sequences of these four merged contigs were used as reference sequences and reads from each individual library were mapped to these reference sequences using Bowtie2 (66). Consensus sequences were extracted from the library-specific contigs. To obtain more of the genome sequence of the lagovirus detected in MF-150, reads and contigs assembled from the MF-150 duodenum library and the Mulligan’s Flat library containing MF-150 duodenum (MF3-D) were aligned to an EBHSV reference sequence (KC832839.1/EBHSV/SWE/O4021-9/1982) using the Geneious mapper tool (68). Two contigs and six reads aligned to EBHSV in regions upstream of the MF-150 Sanger sequence already obtained, allowing the design of primers to amplify and sequence across missing regions.

#### Genome confirmation and extension PCRs

Following RNA sequencing, further primers were designed to amplify missing parts of the newly identified hare calicivirus genomes, as well as to confirm the genome sequence for HaCV-A1 (JM-29 duodenum) by amplicon sequencing (Supplementary table S1). Primer GV271 (62) was used for 3’ end amplification from within the polyA tail. PCRs were conducted using Invitrogen Platinum *Taq* Polymerase High Fidelity kit according to the manufacturer’s protocol, using 2.4 μ1 of cDNA as template in a 40 μ1 reaction. DNA libraries were prepared and sequenced using either Illumina Miseq technology as described previously (61, 69) or Sanger sequencing conducted at ACRF-BRF.

Consensus sequences for the near complete genome of HaCV-A1, HaCV-A2 and partial genome of HaCV-A3 have been deposited in GenBank under accession numbers MK138383-MK138385. All RNA-Seq reads were deposited into the NCBI sequence read archive (SRA) under BioProject XXXX.

#### Identification of conserved domains and potential ORFs

The NCBI Conserved domains tool (70) was used to check for the presence of conserved functional domains in the newly discovered complete and partial viral genomes, and the ExPASY translate tool (https://web.expasy.org/translate/) and the Geneious ORF prediction tool were used to identify realistic open reading frames. Although several possible ORFs existed, the location of ORF 1 in all three viruses was inferred due to its size (largest ORF) and through sequence similarity with other lagoviruses (accession NC_002615, NC_001543). The location of ORF 2 (in HaCV-A1 and HaCV-A2) was chosen based on the location of putative TURBS motifs (which were found manually), as translation re-initiation in caliciviruses tends to occur within 12 – 24 nt of the TURBS structure, and only one potential ORF fit this criteria (71). The Geneious annotate and predict tool was used to annotate the genomes based on published lagovirus sequences, and the likely cleavage fragments of the ORF 1 polyprotein were inferred from sequence homology with EBHSV and RFIDV (accession NC_002615, NC_001543).

### Recombination analyses

RDP4 (60) was used to screen for evidence of recombination with a data set containing the three new sequences plus 18 non-recombinant lagovirus sequences, including both rabbit and hare viruses. The data set was trimmed to the length of the HaCV-A3 sequence (alignment length 4,588 nt). The RDP, GENECONV and MAXCHI methods were used to explore data for recombination signals, and BOOTSCAN and CHIMAERA were used to verify signals detected by initial screening methods. A p-value of 0.05 represented a significant result for all tests, and putative recombination events were considered to be those detected by at least two of the three initial methods. A pairwise identity plot was generated by the RDP method with a sliding window of 30 nt. To confirm recombination events, we inferred phylogenetic trees on sections of the alignment either side of the putative recombination break-point using a maximum likelihood approach as described below. Significant evidence for recombination was reported as cases of clear phylogenetic incongruence with strong (i.e. >70%) bootstrap support.

### Phylogenetic analysis

The lagovirus genome sequences identified here were aligned with 28 (24 complete genomes, four capsid sequence only) sequences available on GenBank, representing the known diversity of lagoviruses, using MAFFT as available in Geneious (68). Maximum likelihood (ML) phylogenetic trees were inferred for both the RdRp (1,548 nt, 27 sequences) and capsid (1,704 nt, 31 sequences) using PhyML (72) and employing the GTR+Γ+I model of nucleotide substitution (as selected using jModelTest V2.1.6 (73,74)) with five rate categories, an estimated proportion of invariant sites and gamma distribution parameter. Topology searching used a combination of nearest-neighbor interchange and subtree pruning and regrafting branch-swapping. Branch support was estimated using 1,000 bootstrap replicates using the same ML procedure as described above, and all trees were mid-point rooted for clarity.

### Hare calicivirus screening PCRs

To determine the prevalence of each of the new hare caliciviruses, hare duodenum RNA was individually screened for each new lagovirus. Specific primer sets were designed based on the sequence of the three new lagoviruses to enable detection of each virus; HaCV-A1, HareCaV1_F6.2 and HareCaV1_R6.5 (330 bp amplicon); HaCV-A2, HareCaV2_F5.2 and HareCaV2_R5.4 (213 bp amplicon); HaCV-A3, HareCaV4_F5.5 and HareCaV4_R5.g (408 bp amplicon) (Supplementary table S1). Liver RNA was also screened from hares for which a product was amplified from the duodenum RNA. RT-PCR was conducted using the One-Step Ahead RT-PCR kit (Qiagen, Hilden, Germany) according to the manufacturer’s instructions using 1 μ1 of RNA diluted 1:10 in nuclease free water in a 10 μ1 reaction volume.

## Acknowledgements

ECH is supported by an ARC Australian Laureate Fellowship (FL170100022). We thank John Matthews and Alex Thorpe from Agriculture Victoria as well as Oliver Orgill and team from the ACT Parks and Conservation Services for sample acquisition.

Data available at the NCBI sequence read archive (SRA) BioProject XXXX, and in GenBank under accession numbers MK138383-MK138385.

## References

1. Koonin EV, Dolja VV, Krupovic M. 2015. Origins and evolution of viruses of eukaryotes: The ultimate modularity. Virology 479-480:2-25.

2. Zhang Y-Z, Shi M, Holmes EC. 2018. Using Metagenomics to Characterize an Expanding Virosphere. Cell 172:1168-1172.

3. Shi M, Zhang YZ, Holmes EC. 2018. Meta-transcriptomics and the evolutionary biology of RNA viruses. Virus Res 243:83-90.

4. Shi M, Lin XD, Chen X, Tian JH, Chen LJ, Li K, Wang W, Eden JS, Shen JJ, Liu L, Holmes EC, Zhang YZ. 2018. The evolutionary history of vertebrate RNA viruses. Nature 556:197-202.

5. Shi M, Lin XD, Tian JH, Chen LJ, Chen X, Li CX, Qin XC, Li J, Cao JP, Eden JS, Buchmann J, Wang W, Xu J, Holmes EC, Zhang YZ. 2016. Redefining the invertebrate RNA virosphere. Nature 540:539-543.

6. Li CX, Shi M, Tian JH, Lin XD, Kang YJ, Chen LJ, Qin XC, Xu J, Holmes EC, Zhang YZ. 2015. Unprecedented genomic diversity of RNA viruses in arthropods reveals the ancestry of negative-sense RNA viruses. Elife 4.

7. Calisher CH, Childs JE, Field HE, Holmes KV, Schountz T. 2006. Bats: important reservoir hosts of emerging viruses. Clin Microbiol Rev 19:531-45.

8. Domingo E. 2010. Mechanisms of viral emergence. Vet Res 41:38.

9. Watson DC, Sargianou M, Papa A, Chra P, Starakis I, Panos G. 2014. Epidemiology of Hantavirus infections in humans: a comprehensive, global overview. Crit Rev Microbiol 40:261-72.

10. Geoghegan JL, Holmes EC. 2018. The phylogenomics of evolving virus virulence. Nat Rev Genet doi:10.1038/s41576-018-0055-5.

11. Capucci L, Fusi P, Lavazza A, Pacciarini ML, Rossi C. 1996. Detection and preliminary characterization of a new rabbit calicivirus related to rabbit hemorrhagic disease virus but nonpathogenic. J Virol 70:8614-23.

12. Le Gall-Recule G, Zwingelstein F, Fages MP, Bertagnoli S, Gelfi J, Aubineau J, Roobrouck A, Botti G, Lavazza A, Marchandeau S. 2011. Characterisation of a non-pathogenic and non-protective infectious rabbit lagovirus related to RHDV. Virology 410:395-402.

13. Le Pendu J, Abrantes J, Bertagnoli S, Guitton JS, Le Gall-Recule G, Lopes AM, Marchandeau S, Alda F, Almeida T, Celio AP, Barcena J, Burmakina G, Blanco E, Calvete C, Cavadini P, Cooke B, Dalton K, Delibes Mateos M, Deptula W, Eden JS, Wang F, Ferreira CC, Ferreira P, Foronda P, Goncalves D, Gavier-Widen D, Hall R, Hukowska-Szematowicz B, Kerr P, Kovaliski J, Lavazza A, Mahar J, Malogolovkin A, Marques RM, Marques S, Martin-Alonso A, Monterroso P, Moreno S, Mutze G, Neimanis A, Niedzwiedzka-Rystwej P, Peacock D, Parra F, Rocchi M, Rouco C, Ruvoen-Clouet N, Silva E, Silverio D, Strive T, Thompson G, et al. 2017. Proposal for a unified classification system and nomenclature of lagoviruses. J Gen Virol 98:1658-1666.

14. Strive T, Wright JD, Robinson AJ. 2009. Identification and partial characterisation of a new Lagovirus in Australian wild rabbits. Virology 384:97-105.

15. Cavadini P, Molinari S, Pezzoni G, Chiari M, Brocchi E, Lavazza A, Capucci L. 2016. Identification of a new non-pathogenic lagovirus in brown hares (*Lepus europeaus*), p. In Kelly P, Phillips S, Smith A, Browning C (ed), 5th World Lagomorph Conference. Turlock, CA: California State University Stanislaus.

16. Droillard C, Lemaitre E, Chatel M, Guitton J-S, Marchandeau S, Eterradossi N, Le Gall-Recule G. First complete genome sequence of hare calicivirus isolated from Lepus europaeus. MRA, in press.

17. Wirblich C, Meyers G, Ohlinger VF, Capucci L, Eskens U, Haas B, Thiel HJ. 1994. European brown hare syndrome virus: relationship to rabbit hemorrhagic disease virus and other caliciviruses. J Virol 68:5164-73.

18. Abrantes J, van der Loo W, Le Pendu J, Esteves PJ. 2012. Rabbit haemorrhagic disease (RHD) and rabbit haemorrhagic disease virus (RHDV): a review. Vet Res 43:12.

19. Le Gall G, Huguet S, Vende P, Vautherot JF, Rasschaert D. 1996. European brown hare syndrome virus: molecular cloning and sequencing of the genome. J Gen Virol 77 (Pt 8):1693-7.

20. Wirblich C, Thiel HJ, Meyers G. 1996. Genetic map of the calicivirus rabbit hemorrhagic disease virus as deduced from in vitro translation studies. J Virol 70: 7974-83.

21. Meyers G, Wirblich C, Thiel HJ. 1991. Genomic and subgenomic RNAs of rabbit hemorrhagic disease virus are both protein-linked and packaged into particles. Virology 184:677-86.

22. Bull RA, Hansman GS, Clancy LE, Tanaka MM, Rawlinson WD, White PA. 2005. Norovirus recombination in ORF1/ORF2 overlap. Emerg Infect Dis 11:1079-85.

23. Coyne KP, Reed FC, Porter CJ, Dawson S, Gaskell RM, Radford AD. 2006. Recombination of Feline calicivirus within an endemically infected cat colony. J Gen Virol 87:921-926.

24. Clarke IN, Lambden PR. 1997. The molecular biology of caliciviruses. J Gen Virol 78 (Pt 2):291-301.

25. Fuchs A, Weissenbock H. 1992. Comparative histopathological study of rabbit haemorrhagic disease (RHD) and European brown hare syndrome (EBHS). J Comp Pathol 107:103-13.

26. Stott P. 2015. Factors influencing the importation and establishment in Australia of the European hare (*Lepus europaeus*). Aust J Zool 63:46-75.

27. Fenner F. 2010. Deliberate introduction of the European rabbit, *Oryctolagus cuniculus*, into Australia. Rev Sci Tech 29:103-11.

28. Rolls EC. 1984. They All Ran Wild: The Animals and Plants that Plague Australia. Angus & Robertson.

29. Myers K, Parker BS. 1965. A study of the biology of the wild rabbit in climatically different regions in eastern Australia. I. Patterns of distribution. Wildlife Research 10:1-32.

30. Agriculture Victoria. 31 July 2017. Pest Animals, European Hare, on Victoria State Government. http://agriculture.vic.aov.au/agriculture/pests-diseases-and-weeds/pest-animals/a-z-of-pest-animals/european-hare. Accessed 18 September 2018.

31. Government of South Australia. 2017. Consolidated list of declarations of animals and plants. Department of primary industries, South Australia. http://www.pir.sa.gov.au/_data/assets/pdf_file/0002/221924/Declaration_of_Animals_and_Plants_-_July_2017.pdf.

32. Cooke BD, Fenner F. 2002. Rabbit haemorrhagic disease and the biological control of wild rabbits, *Oryctolagus cuniculus*, in Australia and New Zealand. Wildlife Research 29:689-706.

33. Nagesha HS, McColl KA, Collins BJ, Morrissy CJ, Wang LF, Westbury HA. 2000. The presence of cross-reactive antibodies to rabbit haemorrhagic disease virus in Australian wild rabbits prior to the escape of virus from quarantine. Arch Virol 145: 749-57.

34. Strive T, Elsworth P, Liu J, Wright JD, Kovaliski J, Capucci L. 2013. The non-pathogenic Australian rabbit calicivirus RCV-A1 provides temporal and partial cross protection to lethal Rabbit Haemorrhagic Disease Virus infection which is not dependent on antibody titres. Vet Res 44:51.

35. Strive T, Wright J, Kovaliski J, Botti G, Capucci L. 2010. The non-pathogenic Australian lagovirus RCV-A1 causes a prolonged infection and elicits partial cross-protection to rabbit haemorrhagic disease virus. Virology 398:125-34.

36. Lemaitre E, Zwingelstein F, Marchandeau S, Le Gall-Recule G. 2018. First complete genome sequence of a European non-pathogenic rabbit calicivirus (lagovirus GI.3). Arch Virol 163:2921-2924.

37. Nicholson LJ, Mahar JE, Strive T, Zheng T, Holmes EC, Ward VK, Duckworth JA. 2017. Benign Rabbit Calicivirus in New Zealand. Appl Environ Microbiol 83.

38. Mahar JE, Read AJ, Gu X, Urakova N, Mourant R, Piper M, Haboury S, Holmes EC, Strive T, Hall RN. 2018. Detection and Circulation of a Novel Rabbit Hemorrhagic Disease Virus in Australia. Emerg Infect Dis 24:22-31.

39. Hall RN, Mahar JE, Haboury S, Stevens V, Holmes EC, Strive T. 2015. Emerging Rabbit Hemorrhagic Disease Virus 2 (RHDVb), Australia. Emerg Infect Dis 21:2276-8.

40. Hall RN, Mahar JE, Read AJ, Mourant R, Piper M, Huang N, Strive T. 2018. A strain-specific multiplex RT-PCR for Australian rabbit haemorrhagic disease viruses uncovers a new recombinant virus variant in rabbits and hares. Transbound Emerg Dis 65:e444-e456.

41. Hall RN, Peacock DE, Kovaliski J, Mahar JE, Mourant R, Piper M, Strive T. 2017. Detection of RHDV2 in European brown hares (*Lepus europaeus*) in Australia. Vet Rec 180:121.

42. Mahar JE, Hall RN, Peacock D, Kovaliski J, Piper M, Mourant R, Huang N, Campbell S, Gu X, Read A, Urakova N, Cox T, Holmes EC, Strive T. 2017. Rabbit haemorrhagic disease virus 2 (GI.2) is replacing endemic strains of RHDV in the Australian landscape within 18 months of its arrival. J Virol doi:10.1128/JVI.01374-17.

43. Duff JP, Chasey D, Munro R, Wooldridge M. 1994. European brown hare syndrome in England. Vet Rec 134:669-73.

44. Frolich K, Fickel J, Ludwig A, Lieckfeldt D, Streich WJ, Jurcik R, Slamecka J, Wibbelt G. 2007. New variants of European brown hare syndrome virus strains in free-ranging European brown hares (*Lepus europaeus*) from Slovakia. J Wildl Dis 43:89-96.

45. Billinis C, Psychas V, Tontis DK, Spyrou V, Birtsas PK, Sofia M, Likotrafitis F, Maslarinou OM, Kanteres D. 2005. European brown hare syndrome in wild European brown hares from Greece. J Wildl Dis 41: 783-6.

46. Syrjala P, Nylund M, Heinikainen S. 2005. European brown hare syndrome in free-living mountain hares (*Lepus timidus*) and European brown hares (*Lepus europaeus*) in Finland 1990-2002. J Wildl Dis 41:42-7.

47. Frolich K, Haerer G, Bacciarini L, Janovsky M, Rudolph M, Giacometti M. 2001. European brown hare syndrome in free-ranging European brown and mountain hares from Switzerland. J Wildl Dis 37:803-7.

48. Le Gall-Recule G, Zwingelstein F, Laurent S, Portejoie Y, Rasschaert D. 2006. Molecular epidemiology of European brown hare syndrome virus in France between 1989 and 2003. Arch Virol 151:1713-21.

49. Le Gall-Recule G, Lemaitre E, Bertagnoli S, Hubert C, Top S, Decors A, Marchandeau S, Guitton JS. 2017. Large-scale lagovirus disease outbreaks in European brown hares (*Lepus europaeus*) in France caused by RHDV2 strains spatially shared with rabbits (*Oryctolagus cuniculus*). Vet Res 48: 70.

50. Velarde R, Cavadini P, Neimanis A, Cabezon O, Chiari M, Gaffuri A, Lavin S, Grilli G, Gavier-Widen D, Lavazza A, Capucci L. 2017. Spillover Events of Infection of Brown Hares (*Lepus europaeus*) with Rabbit Haemorrhagic Disease Type 2 Virus (RHDV2) Caused Sporadic Cases of an European Brown Hare Syndrome-Like Disease in Italy and Spain. Transbound Emerg Dis 64:1750-1761.

51. Camarda A, Pugliese N, Cavadini P, Circella E, Capucci L, Caroli A, Legretto M, Mallia E, Lavazza A. 2014. Detection of the new emerging rabbit haemorrhagic disease type 2 virus (RHDV2) in Sicily from rabbit (Oryctolagus cuniculus) and Italian hare (Lepus corsicanus). Res Vet Sci 97:642-5.

52. Puggioni G, Cavadini P, Maestrale C, Scivoli R, Botti G, Ligios C, Le Gall-Recule G, Lavazza A, Capucci L. 2013. The new French 2010 Rabbit Hemorrhagic Disease Virus causes an RHD-like disease in the Sardinian Cape hare (Lepus capensis mediterraneus). Vet Res 44:96.

53. Royall E, Locker N. 2016. Translational Control during Calicivirus Infection. Viruses 8:104.

54. Lopes AM, Capucci L, Gavier-Widen D, Le Gall-Recule G, Brocchi E, Barbieri I, Quemener A, Le Pendu J, Geoghegan JL, Holmes EC, Esteves PJ, Abrantes J. 2014. Molecular evolution and antigenic variation of European brown hare syndrome virus (EBHSV). Virology 468-470:104-12.

55. Liu J, Fordham DA, Cooke BD, Cox T, Mutze G, Strive T. 2014. Distribution and prevalence of the Australian non-pathogenic rabbit calicivirus is correlated with rainfall and temperature. PLoS One 9:e113976.

56. Chasey D, Duff P. 1990. European brown hare syndrome and associated virus particles in the UK. Vet Rec 126:623-4.

57. Le Gall-Reculé G, Lavazza A, Marchandeau S, Bertagnoli S, Zwingelstein F, Cavadini P, Martinelli N, Lombardi G, Guérin J-L, Lemaitre E, Decors A, Boucher S, Le Normand B, Capucci L. 2013. Emergence of a new lagovirus related to Rabbit Haemorrhagic Disease Virus. Vet Res 44:81.

58. Lopes AM, Dalton KP, Magalhaes MJ, Parra F, Esteves PJ, Holmes EC, Abrantes J. 2015. Full genomic analysis of new variant rabbit hemorrhagic disease virus (RHDVb) revealed multiple recombination events. J Gen Virol 96(Pt 6):1309-19.

59. Lopes AM, Dalton KP, Magalhaes MJ, Parra F, Esteves PJ, Holmes EC, Abrantes J. 2015. Full genomic analysis of new variant rabbit hemorrhagic disease virus revealed multiple recombination events. J Gen Virol 96:1309-19.

60. Martin DP, Murrell B, Golden M, Khoosal A, Muhire B. 2015. RDP4: Detection and analysis of recombination patterns in virus genomes. Virus Evol 26.

61. Mahar JE, Nicholson L, Eden JS, Duchene S, Kerr PJ, Duckworth J, Ward VK, Holmes EC, Strive T. 2016. Benign Rabbit Caliciviruses Exhibit Evolutionary Dynamics Similar to Those of Their Virulent Relatives. J Virol 90:9317-29.

62. Eden JS, Tanaka MM, Boni MF, Rawlinson WD, White PA. 2013. Recombination within the pandemic norovirus GII.4 lineage. J Virol 87:6270-82.

63. Bolger AM, Lohse M, Usadel B. 2014. Trimmomatic: a flexible trimmer for Illumina sequence data. Bioinformatics 30:2114-20.

64. Grabherr MG, Haas BJ, Yassour M, Levin JZ, Thompson DA, Amit I, Adiconis X, Fan L, Raychowdhury R, Zeng Q, Chen Z, Mauceli E, Hacohen N, Gnirke A, Rhind N, di Palma F, Birren BW, Nusbaum C, Lindblad-Toh K, Friedman N, Regev A. 2011. Full-length transcriptome assembly from RNA-Seq data without a reference genome. Nat Biotechnol 29:644-52.

65. Li B, Ruotti V, Stewart RM, Thomson JA, Dewey CN. 2010. RNA-Seq gene expression estimation with read mapping uncertainty. Bioinformatics 26:493-500.

66. Langmead B, Salzberg SL. 2012. Fast gapped-read alignment with Bowtie 2. Nat Methods 9:357-9.

67. Quast C, Pruesse E, Yilmaz P, Gerken J, Schweer T, Yarza P, Peplies J, Glockner FO. 2013. The SILVA ribosomal RNA gene database project: improved data processing and web-based tools. Nucleic Acids Res 41:D590-6.

68. Kearse M, Moir R, Wilson A, Stones-Havas S, Cheung M, Sturrock S, Buxton S, Cooper A, Markowitz S, Duran C, Thierer T, Ashton B, Meintjes P, Drummond A. 2012. Geneious Basic: an integrated and extendable desktop software platform for the organization and analysis of sequence data. Bioinformatics 28:1647-9.

69. Eden JS, Kovaliski J, Duckworth JA, Swain G, Mahar JE, Strive T, Holmes EC. 2015. Comparative Phylodynamics of Rabbit Hemorrhagic Disease Virus in Australia and New Zealand. J Virol 89:9548-58.

70. Marchler-Bauer A, Bo Y, Han L, He J, Lanczycki CJ, Lu S, Chitsaz F, Derbyshire MK, Geer RC, Gonzales NR, Gwadz M, Hurwitz DI, Lu F, Marchler GH, Song JS, Thanki N, Wang Z, Yamashita RA, Zhang D, Zheng C, Geer LY, Bryant SH. 2017. CDD/SPARCLE: functional classification of proteins via subfamily domain architectures. Nucleic Acids Res 45:D200-d203.

71. Luttermann C, Meyers G. 2014. Two alternative ways of start site selection in human norovirus reinitiation of translation. J Biol Chem 289:11739-54.

72. Guindon S, Dufayard JF, Lefort V, Anisimova M, Hordijk W, Gascuel O. 2010. New algorithms and methods to estimate maximum-likelihood phylogenies: assessing the performance of PhyML 3.0. Syst Biol 59:307-21.

73. Darriba D, Taboada GL, Doallo R, Posada D. 2012. jModelTest 2: more models, new heuristics and parallel computing. Nat Methods 9: 772.

74. Guindon S, Gascuel O. 2003. A simple, fast, and accurate algorithm to estimate large phylogenies by maximum likelihood. Syst Biol 52:696-704.

